# The Role of UBE2-Conjugating Enzymes in the Mechanism of MuRF1 Ubiquitylation

**DOI:** 10.1101/2022.06.13.495268

**Authors:** Peter W.J. Dawson, Leslie M. Baehr, David C. Hughes, Timothy J. Knowles, Pooja Sridhar, Sue C. Bodine, Yu-Chiang Lai

## Abstract

MuRF1 (Muscle-specific RING finger protein 1; gene name TRIM63) is a ubiquitin E3 ligase, associated with the progression of muscle atrophy. As a RING (Really Interesting New Gene)-type E3 ligase, its unique activity of ubiquitylation is driven by a specific interaction with UBE2 (ubiquitin conjugating enzyme) to ubiquitylate its substrate protein. The understanding of MuRF1 function remains unclear as candidate UBE2 has not been elucidated and thus the mechanism of ubiquitylation is inconclusive. In the present study, we screened human ubiquitin dependent E2s using in-vitro ubiquitylation assays. We found that MuRF1 engages in ubiquitylation/auto-ubiquitylation with UBE2D, UBE2E, UBE2N/V families and UBE2W. Our result indicated that MuRF1 can cause mono-ubiquitylation, K48, and K63 specific poly-ubiquitin chains in a UBE2-dependent manner. Interestingly, we identified a two-step UBE2-dependent mechanism by which UBE2W allows MuRF1 to mono-ubiquitylate which then acts as an anchor for UBE2N/V and UBE2D generated poly-ubiquitin chain formation. Furthermore, MuRF1 was shown to cooperate with the identified interacting UBE2s to directly ubiquitylate substrates Titin (A168-A170), Desmin, and MYLPF (Myosin Light Chain, Phosphorylatable, Fast Skeletal Muscle; also called Myosin Light Regulatory Chain 2). Our work presents a novel insight into the mechanisms that underpin MuRF1 activity by highlighting the diversity of MuRF1 ubiquitylation enabled by different UBE2s.

## INTRODUCTION

MuRF1, a muscle-specific TRIM (tripartite motif-containing) E3 ligase (TRIM63), plays a critical role that causes skeletal muscle atrophy. Many studies, including in humans and mice, demonstrate that the expression of the MuRF1 gene (mRNA) and protein expressions are increased following disuse, heart failure, lung injury, fasting, cancer, renal failure, diabetes, cytokine exposure, corticosteroid treatment and denervation [1–7]. Importantly, suppression of MuRF1 expression perturbs atrophy induced by glucocorticoid treatment, limb unloading, denervation, and lung injury [1,7–11]. These earlier studies have established MuRF1 as a key regulator of muscle mass. Nonetheless, insight into the molecular mechanisms of MuRF1-induced muscle atrophy are unclear, slowing the development of therapeutics that target MuRF1-induced muscle atrophy.

As MuRF1’s function as a E3 ligase is to ubiquitylate a substrate, many potential candidate substrates have been reported [8,11–23]. However, only few substrates have been shown to be directly ubiquitylated and subsequently degraded. The process of protein ubiquitylation is a coordinated sequence of three enzymatic actions: by an E1 ubiquitin-activating enzyme (UBE1), an E2 ubiquitin-conjugating enzyme (UBE2), and finally an E3 ubiquitin-ligase. The UBE1 enzyme hydrolyses ATP to adenylate ubiquitin, which is transferred to an UBE2 active site. As MuRF1 is a RING-type (Really Interesting New Gene) E3 ligase, with no catalytic activity, it requires the interaction of an UBE2 and substrate simultaneously to directly transfer ubiquitin from UBE2 to MuRF1 substrate(s). UBE2s are responsible for the type of ubiquitylation (Mono- or Poly-ubiquitylation) and the structure of ubiquitin chains based on ubiquitin-ubiquitin lysine attachment [24,25]. There are eight different lysine-conjugated ubiquitin chain types (M1, K6, K11, K27, K29, K33, K48 and K63) and the topology of ubiquitin chain types ultimately determine the fate of the target protein, such as degradation, delocalisation, or other signalling events [26]. Therefore, to understand the functional role of MuRF1 ubiquitylation onto its substrate, one must identify the UBE2 that partners with MuRF1. Identifying MuRF1 partnering E2s also provide a tool to directly explore substrates of MuRF1 *in-vitro* and characterise their specific form of ubiquitylation.

Previous studies have attempted to identify UBE2s that interact with MuRF1, with no measurement of ubiquitylation activity. Polge *et al*. (2018) applied yeast two-hybrid screen and SPR (surface plasmon resonance) technologies and identified several UBE2s, including UBE2E1, UBE2G1, UBE2J1, UBE2J2, and UBE2L3, as interacting with MuRF1. However, these two methods only measure protein-protein interaction without detecting ubiquitin E3 ligase activity of MuRF1. When MuRF1-UBE2 ubiquitylation activity has been studied using ELISA based methods, only 11 UBE2s have been explored [28]. Further studies that have measured MuRF1 ubiquitylation activity using in-vitro assays, only use UBE2D3 (UBCH5C) as an UBE2-interactor [13,17,20,29]. While this offers some insights into MuRF1-UBE2D3 ubiquitylation activity, it is worth noting that the UBE2D family may be promiscuous and can interact and produce ubiquitylation activity with most RING type E3 ligases [25]. The current literature offers limited understanding of MuRF1-UBE2 interactors and how they relate to MuRF1 ubiquitylation function. Therefore, a study of all human UBE2s with MuRF1 ubiquitylation is necessary to further understand the mechanism of MuRF1-mediated ubiquitylation.

In the present study a full human UBE2 library (excluding ubiquitin-like UBE2s; see table S1) were screened using a standard in-vitro ubiquitylation assay. In addition to the UBE2D and UBE2E families, the present study showed the UBE2N/V family and UBE2W also interacting with MuRF1. Moreover, MuRF1 was also found to interact with UBE2W and UBE2N/Vs in a sequential two-step fashion to ubiquitylate substrates and forming K63-specific ubiquitin chain. This UBE2-MuRF1 function was found to be consistent with both auto-ubiquitylation and ubiquitylation of MuRF1 substrates, Titin, Desmin, and MYLPF.

## RESULTS

### MuRF1 interacts with UBE2D, E, N/V families, and UBE2W to form poly-ubiquitin chains when using cy-5 tagged fluorescent ubiquitin

To investigate which E2s interact with MuRF1, a full screen of 28 recombinant human UBE2 enzymes was undertaken to determine which catalyse MuRF1-dependent ubiquitylation (Fig 1). *In-vitro* ubiquitylation assays were performed using cy-5 labelled ubiquitin as a substrate. The results showed that MuRF1 interacts with UBE2D family (including UBE2D1, D2, D3 and D4), UBE2E family (including UBE2E1, E2, and E3), UBE2N/V1 & N/V2 and UBE2W by forming poly-ubiquitin chains in presence of MBP-MuRF1. MBP-MuRF1 (anti-MBP blot) was revealed to be modified, indicating auto-ubiquitylation. Coomassie blue gel staining was performed to show the presence of each protein loaded in the assay (Fig 1).

**Figure 1.**
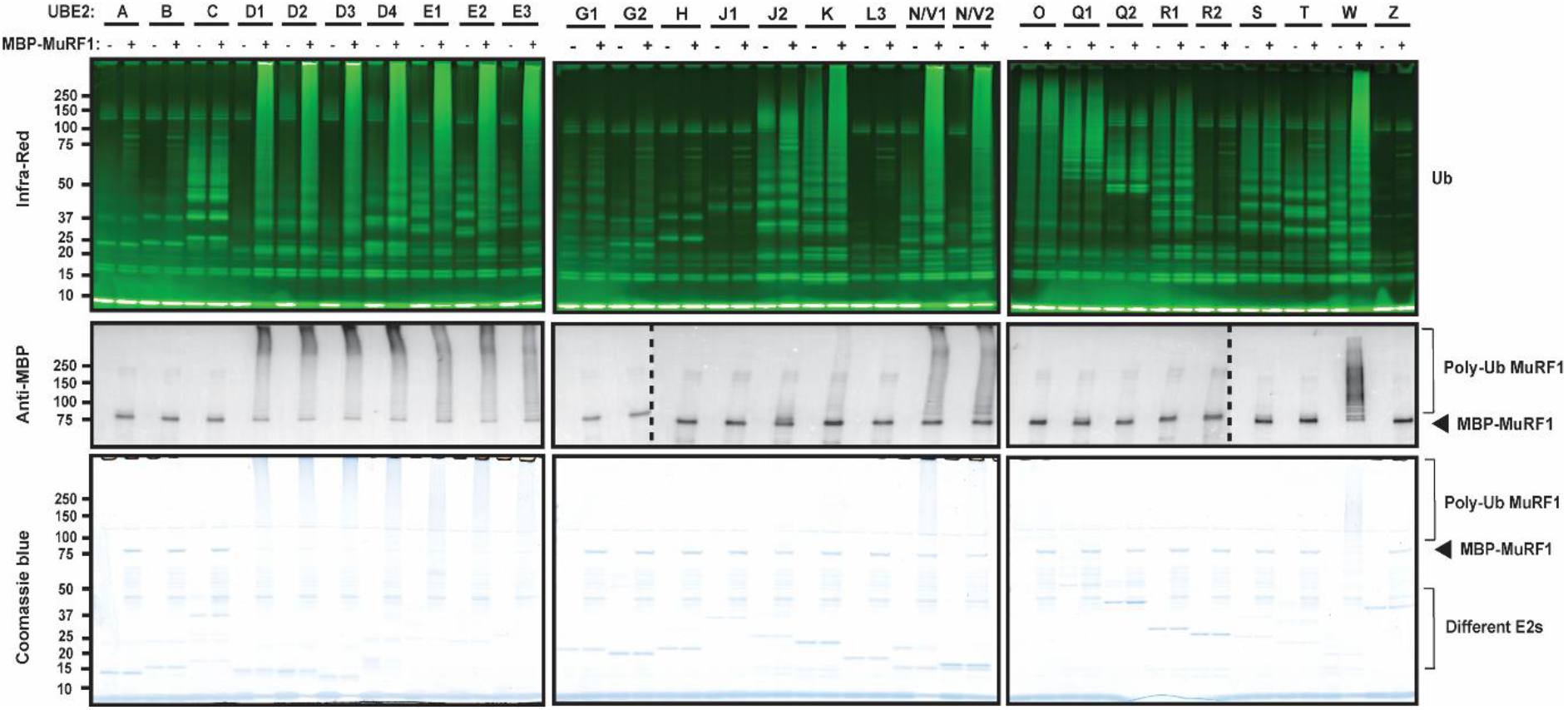
Ubiquitylation assay, using Cy-5 labelled ubiquitin, shows UBE2D, UBE2E, UBE2 N/V, and UBE2W family members partnering with MuRF1 to form poly-ubiquitin chains. A library of 28 UBE2s (With exception to non-classical Ubiquitin E2s - listed in Table S1) were incubated with or without MBP-MuRF1 for ubiquitylation reactions. After incubation for 1 hr, samples were subject to SDS-PAGE to detect ubiquitin chain formation using Li-cor Odyssey CLx to detect (cy-5 tagged) fluorescent ubiquitin (Upper panel). MuRF1 auto-ubiquitylation was detected by immunoblot using anti-MBP (Middle panel). Sample loadings were visualised using Coomassie blue staining (Lower panel).

### MuRF1 produces distinct ubiquitylation (auto-, mono-, poly-, and unanchored-ubiquitylation) in an E2 dependent manner

To confirm the findings from the UBE2 screening, a smaller panel was assayed with native ubiquitin (without cy-5 labelling). We confirmed that UBE2D and UBE2E families partner with MBP-MuRF1 to form poly-ubiquitin chains (Fig 2B) and MuRF1 auto-ubiquitylation (Fig 2A). In contrast to the previous E2 screening, UBE2N/V1 and UBE2N/V2 generated poly-ubiquitin chains (Fig 2B) without ubiquitylating MBP-MuRF1 itself, suggesting UBE2N/V1 & 2 are not involved in auto-ubiquitylation. Furthermore, UBE2W only mono-ubiquitylated MBP-MuRF1 (Fig 2A) but did not make poly-ubiquitin chains (Fig 2B). UBE2 J, K, Q1, and S did not cause any notable change in protein ubiquitylation.

**Figure 2.**
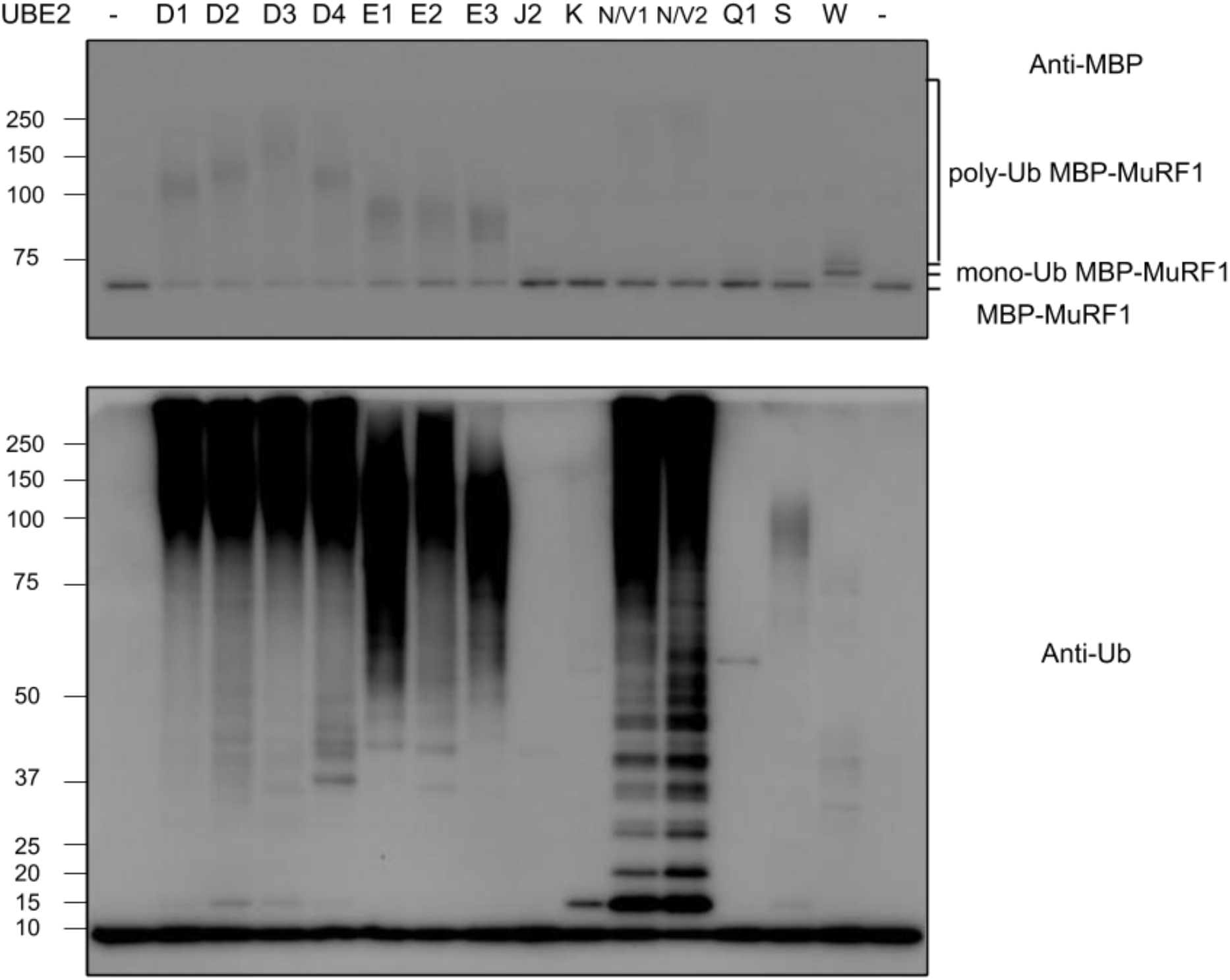
Ubiquitylation assay using native ubiquitin shows UBE2D, E and N/V family partnering with MuRF1 to form poly-ubiquitin chains, whereas UBE2W causes mono-auto-ubiquitylation. Selected ten E2s identified as MuRF1 interactors (Fig.1), plus 4 non-interactors (UBE2J2, K, Q1, and S) as negative controls, were incubated with MBP-MuRF1 for the ubiquitination reactions. After 1 hr incubation, samples were subject to western blotting to detect MuRF1 auto-ubiquitylation using anti-MBP antibody (A) and detect ubiquitin chain formation using anti-ubiquitin antibody (B).

### MuRF1 auto-ubiquitylates by a sequential interaction with UBE2W then UBE2N/V family to form K63 poly-ubiquitin chains

Previous research has shown another TRIM E3-ligase, TRIM5a, partners with UBE2W to generate mono-ubiquitylation as an anchor to attach additional poly-ubiquitin chains [30]. We proceeded to explore if this mechanism occurs with MuRF1. By examining the ubiquitylation generated by each MuRF1-interacting UBE2 in the presence or absence of UBE2W, we found that UBE2N/V1 or UBE2N/V2 alone can only form unanchored poly-ubiquitin chains (Fig 2 & 3B) in the absence of UBE2W. UBE2W alone facilitates the auto-ubiquitylation of MBP-MuRF1 by adding a single ubiquitin (mono-ubiquitylation; Fig 3A). However, the combination of UBE2N/V1 or N/V2 with UBE2W causes the poly-ubiquitylation of MBP-MuRF1 (Fig 3A), suggesting the mono-ubiquitin is an anchor on MuRF1 to attach further poly-ubiquitin chains, generated by UBE2N/V1 or N/V2 (Fig 6). Furthermore, probing for specific poly-ubiquitin chain types demonstrated that UBE2N/V1 and N/V2 can generate K63-linked,but not K48-linked, poly-ubiquitin chains (Fig 3C & D), agreeing with previous studies [30,31].

**Figure 3.**
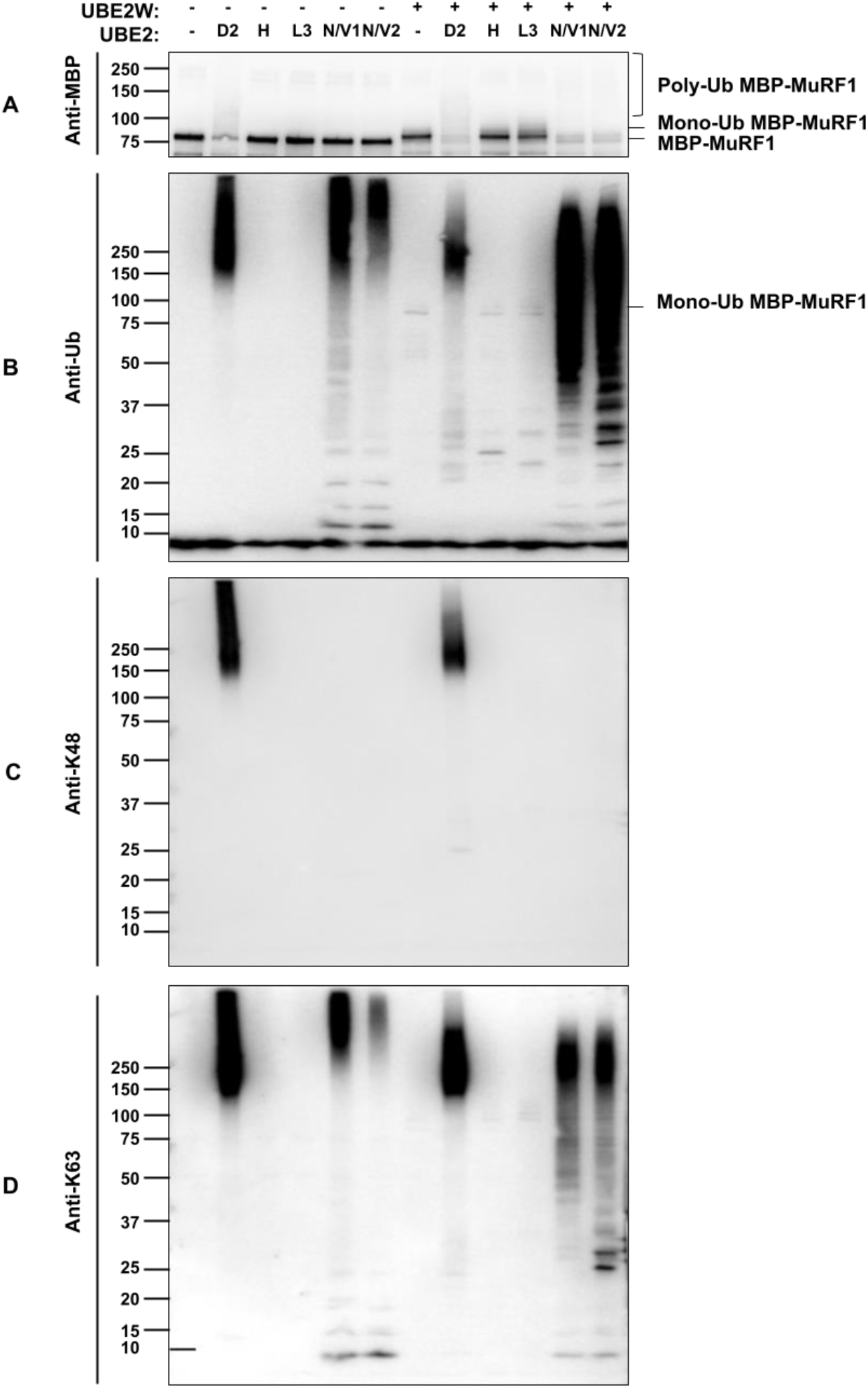
Combination of UBE2W with other MuRF1-interacting E2s shows that UBE2W and UBE2N/V cooperate to generate MuRF1-anchored K63 poly-ubiquitin chains. Ubiquitylation assay of MuRF1-interacting E2s, UBE2D2, N/V1 and N/V2, were incubated in the presence or absence of UBE2W. UBE2H and UBE2L3 were used as negative controls. The reaction mixtures were separated by SDS-PAGE and proteins detected by anti-MBP (A), anti-ubiquitin (B), anti-lysine 48 ubiquitin chains (C) and anti-lysine 63 ubiquitin chains (D).

Consistent with previous experiments (Fig 2), UBE2D2 was able to form poly-ubiquitin chains on MBP-MuRF1. Previous studies have shown UBE2D families are able to generate all eight different linkage types of poly-ubiquitin chains [32] confirmed that UBE2D2 generates both K48- and K63-linked ubiquitin chains (Fig 3C & 3D). However, the addition of UBE2W also modified the poly-ubiquitin chains to a lower molecular weight, suggesting that poly-ubiquitin chains generated by UBE2D2 are also able to attach on the mono-ubiquitin anchor of MBP-MuRF1 (Fig 3A, 3C & 3D).

### MuRF1 directly ubiquitylates previously identified substrates by UBE2W and N/V2 in a two-step mechanism

Moving forward, we explored if MuRF1 ubiquitylation of substrate was the same as MuRF1 auto-ubiquitylation. Titin is well established as a site of MuRF1 translocation and interaction [33,34]. We used a Titin fragment (A168-A170), which has been identified as a MuRF1 interacting domain [35–37], as an *in-vitro* substrate of MuRF1 (Fig 4A). Additionally, we included Desmin and MYLPF identified as potential substrates (Witt *et al*., 2005; Cohen *et al*., 2009; Baehr *et al*., 2021) in our assays. The ubiquitylation assays revealed that MBP-Titin (A168-A170), Desmin, and HIS-MYLPF are ubiquitylated by HIS-SUMO-MuRF1, confirming them to be direct substrates of MuRF1 (Fig 4A). The cooperative mechanism of UBE2W and UBE2N/V enzymes was also seen, with substrates being mono-ubiquitylated by UBE2W, priming them for UBE2N/V2 to form anchored K63 poly-ubiquitin chains onto the substrate (Fig 4A, 4B & 4C). Furthermore, the K48 chain types are exclusive to UBE2D2 interaction and UBE2N/V2 is capable of K63 and not K48 ubiquitin chain formation (Fig 4D).

**Figure 4.**
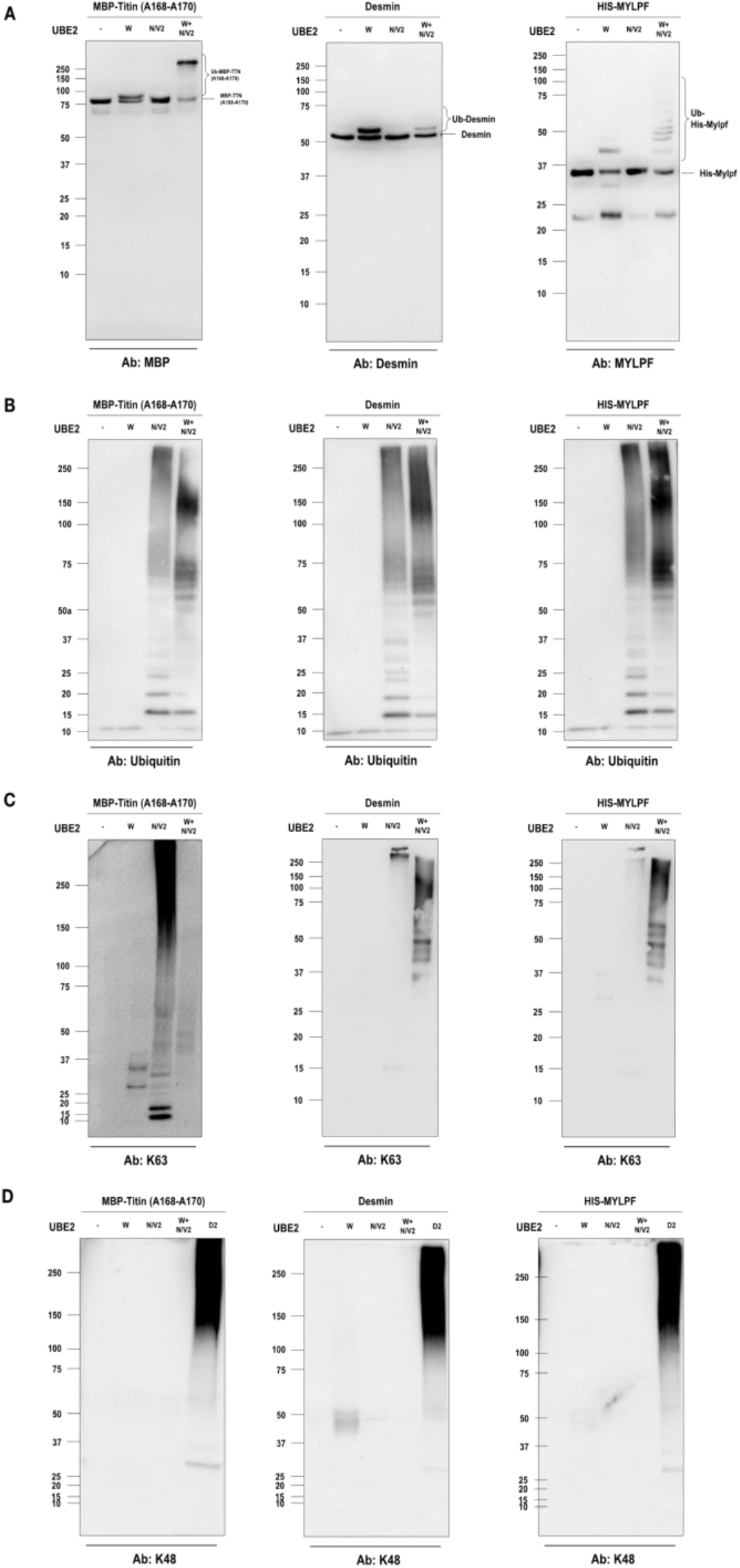
UBE2W and UBE2N/V2 cooperate with MuRF1 to ubiquitylate MBP-Titin (A168-A170), Desmin, and MYLPF substrates through a two-step mechanism. UBE2W and N/V2 were used as UBE2s with HIS-SUMO-MuRF1 and MBP-Titin (A168-A170), Desmin, or HIS-MYLPF and incubated for 1 hr. Proteins were detected using antibodies: anti-MBP, Anti Desmin, Anti-MF-5 (Figure 4A), Anti-ubiquitin (Figure 4B), anti-lysine 63 ubiquitin chains (Figure 4C), and anti-lysine 48 ubiquitin chains (Figure 4D).

### UBE2N, W, and V2 gene expression increases following denervation of mouse skeletal muscle

Having identified MuRF1 cooperating with UBE2W and UBE2N/V to form K63 chains, we wanted to see if this group of enzymes were upregulated in atrophic conditions. Using denervated muscle, that already demonstrated muscle atrophy and increased MuRF1 mRNA expression [38], we assayed for the MuRF1-interacting UBE2s: W, N, V1 and V2. We confirmed the presence of all of these UBE2 in muscle and found that UBE2W, N and V2 are upregulated following 14 days of denervation (Fig 5A, B and D).

**Figure 5.**
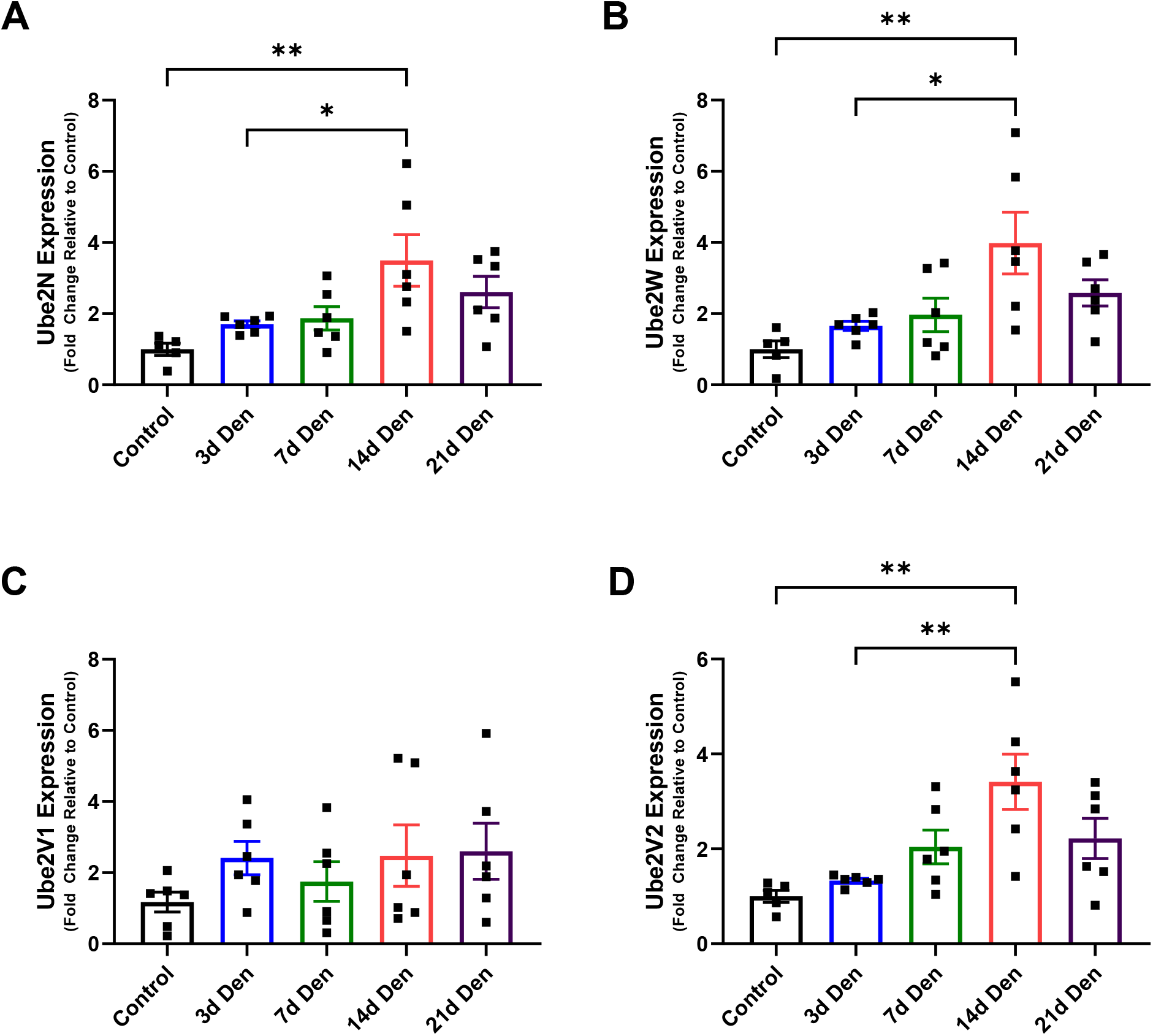
UBE2N, UBE2W, UBE2V2, but not UBE2V1, gene expression increases following denervation of gastrocnemius complex muscles (GSTC) in mice. sC57BL/6 mice were grown for 3-4 months and then had their right leg denervated through surgical ablation of the sciatic nerve. Mice were sacrificed (at days 3, 7, 14, and 21) and their gastrocnemius UBE2 gene expression was measured by rt-qPCR. P-values were calculated using a one-way ANOVA with Tukey’s post hoc test, *P <.05; **P < .01. Data presented as means ± SEM (n = 6).

## DISCUSSION

The in vitro ubiquitylation assay allowed identification of the UBE2s that can partner with MuRF1 and revealed that the UBE2N/V family and UBE2W can cooperate with MuRF1 to ubiquitinate substrates. Further, we showed that MuRF1 partners with different UBE2s to form K48 and K63 poly-ubiquitin chains. We identified that MuRF1 can partner with UBE2W to mono-ubiquitylate itself (auto-ubiquitylation) and substrates, which can then serve as an anchor for K63 poly-ubiquitin chain attachment. Establishing this method, enabled us to identify Titin A168-A170, Desmin, and MYLPF as direct substrates of MuRF1.

We identified that MuRF1 was able to partner with all the UBE2D and E families to form poly-ubiquitin chains. Previous research has already shown the capacity for MuRF1 to interact with UBE2D1, to generate isopeptide linkages with any ubiquitin lysine [39], and our data shows that MuRF1 cooperates with all UBE2D2, 3, & 4 also. Additionally, our data offers clarity over previous work showing the lack of MuRF1 and UBE2D2 interaction (using yeast-two-hybrid and surface plasmon resonance) [40]. We also identified that MuRF1 partners with all UBE2E family, of which UBE2E1 has been reported as a mediator of atrophy in C2C12s during dexamethasone-induced atrophy [14]. Our data highlights MuRF1 ubiquitylation activity through direct cooperation with the UBE2E and D family of enzymes. These two families of UBE2s make up a quarter of the Ubiquitin-Conjugating enzymes in humans, offering a substantial number of partners to facilitate MuRF1 activity.

Our data demonstrates a common familial function of MuRF1 with its cousin TRIM E3s, TRIM21 and TRIM5α (Fig 3 and 4). Like these E3 Ligases, MuRF1 can form K63 poly-ubiquitin chains in a two-step fashion by partnering with UBE2N/V1, or V2 and UBE2W (Fig 6) to form an anchor on the substrate, which can then be used as an attachment point for poly-ubiquitin K63 chains formed by UBE2N/V1 and N/V2 [30,43]. UBE2D2 generated K48 chains were aggregated at a lower molecular weight when combined with UBE2W (Fig 3B), this suggests the same two-step mechanism is applicable with K48 chains. Mono-ubiquitylation has been well explored as a function of UBE2W and specifically it targets the α-amino groups of a protein N-termini [44–46]. This offers an interesting new avenue for MuRF1 mediated ubiquitylation, not only lysine residues, but by targeting N-terminal specific ubiquitylation. In addition, this mono-ubiquitin priming point could potentially act as a substrate for other muscle atrophy-related E3 ligases, e.g. MuRF2, MuRF3, CHIP, MAFBx, TRIM32, TRIM72, and MUSA1 [1,21,39,47]. The possibility of MuRF1-instigated mono-ubiquitylation being used by other E2-E3s pairings, implies a sophisticated complex of ubiquitylation activity. It is therefore crucial to understand the diversity of ubiquitin chain types generated by this mechanism, since chain types determine the fate of substrates [48,49]. Beyond interaction, these MuRF1-interacting UBE2s we observed being upregulated during atrophy. We identified that UBE2N, W, V1 and V2 mRNA expression were present in mouse muscle (Fig 5) and UBE2N, W, and V2 transiently increased at 14 days post-denervation (Fig 5 A, B, D). The UBE2D and E families have been previously shown to increase expression in atrophy; in C2C12 cells, mRNA expression of UBE2D1, 2 and E1 increases following 48 hours of dexamethasone treatment [22,48]. Furthermore, transcriptomics of mice treated with dexamethasone for 14 days have shown increased expression of UBE2D2, D3, N, V1, and V2 [49]. This evidence reveals the treatment and time dependent nature of atrophy-models used. The transient expression of UBE2s shown in our work (Fig 5) highlights the limitation of a using a single time point in previous work [22,49,50]. Having identified MuRF1 partnering UBE2s, future work would benefit from screening multiple time points of atrophy for UBE2s.

**Figure 6.**
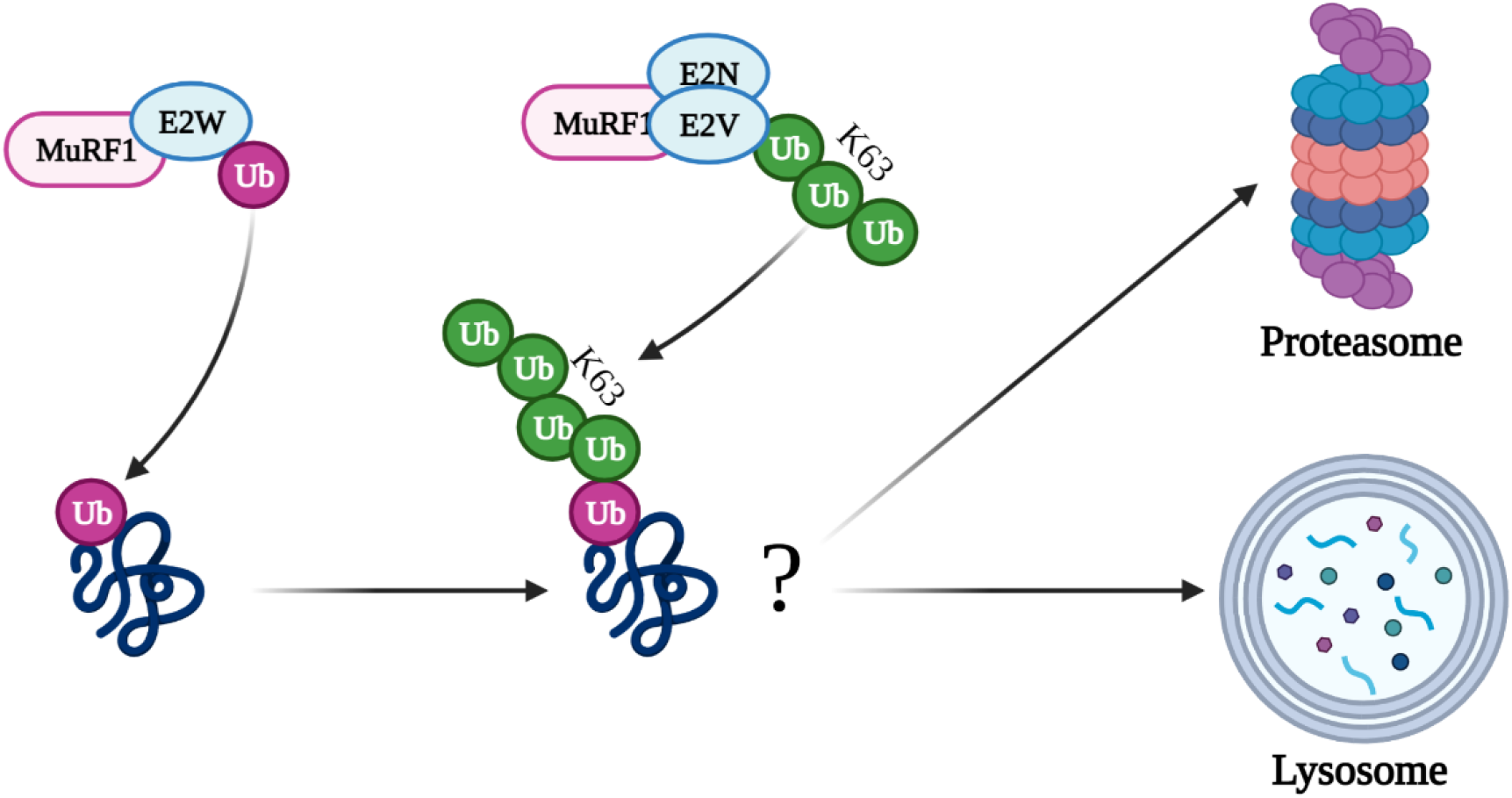
Schematic detailing the two-step mechanism of MuRF1 with UBE2W and UBE2N/V enzymes to generate anchored K63 poly-ubiquitin chains. MuRF1 forms mono-ubiquitylation by cooperating with UBE2W, which acts as an anchor for further ubiquitylation. MuRF1 partnered with UBE2N/V1 or N/V2 form unanchored ubiquitylation, which can utilise the mono-ubiquitin generated by MuRF1 and UBE2W as an anchor. Generated using Biorender.com

Our *in-vitro* data demonstrates that Titin, Desmin and MYLPF are direct substrates of MuRF1 (Fig 4). In agreement with previous studies [37,51], we showed that a fragment of Titin A168-A170 was able to be ubiquitylated by MuRF1 (Fig 4A). We further showed that this ubiquitylation activity is also driven by MuRF1s two-step ubiquitylation with UBE2W and N/V enzymes (Fig 4A). This offers a mechanistic link to previous work showing that Titin is specifically ubiquitylated by K63 chains and MuRF1 knockdown preserves the lysosomal degradation of Titin [52]. In supporting this, a recent study reported that MuRF1 ubiquitylation with the A168-A170 site of Titin resulted in recruitment of NBR1 and P62, proteins that facilitate the autophagy of large protein cargo [51]. These data build a causal chain of events that end with Titin autophagy by p62 and NBR1, mediated by K63 chains, instigated by the MuRF1-E2 interactions that we have elucidated.

Recent screening of the MuRF1 ubiquitylome, using mice overexpressing MuRF1, showed a panel of 56 potential MuRF1 substrates. Further quantification of the potential MuRF1 substrates, by western blot, showed that MYLPF was the only protein decreasing following MuRF1 overexpression, with other protein content remaining stable or increasing [23]. Our data supports previous the finding that MuRF1 is responsible for MYLPF degradation during denervation when comparing wild-type to MuRF1-RING deletion mice [18]. Together this supports the hypothesis that degradation of these substrates is caused directly by MuRF1 ubiquitylation. Desmin has been previously shown to be degraded during atrophy in a TRIM32 dependent manner [45]. Their study showed that while MuRF1 expression is increasing, TRIM32 expression was not increased by fasting-induced atrophy. However, knockdown of TRIM32 prevented the loss of Desmin. Is it possible that MuRF1 and TRIM32 both contribute to Desmin ubiquitylation, by using the mechanism mentioned above, facilitate the degradation of Desmin.

The data presented here offers an insight into the necessary UBE2 enzymes that can enable MuRF1 function. MuRF1 can interact with UBE2D, E, W, and N/V families to ubiquitylate itself and potential substrates. MuRF1 can also generate mono-ubiquitylation and create K63 and K48 poly-ubiquitin chains in a UBE2 dependant manner. Additionally, we presented evidence of direct substrates of MuRF1 ubiquitylation, namely Titin, Desmin, and MYLPF. The further exploration of the role of UBE2W mediated mono-ubiquitylation and its regulation of poly-ubiquitin chains is needed to uncover MuRF1s mechanism of action. Our *in-vitro* findings highlight the context dependent nature of exploring MuRF1 ubiquitylation and stresses the need to examine its interacting partners directly in muscle cells, and preferably during atrophic conditions.

## MATERIALS AND METHODS

### Reagents and antibodies

Recombinant proteins HIS-UBE1, ubiquitin and full library of human UBE2s were sourced from the Medical Research Council - Protein, Phosphorylation and Ubiquitylation Unit (MRC PPU) Reagents and Services (https://mrcppureagents.dundee.ac.uk/). Plasmids for HIS-SUMO-MuRF1 and MBP-MuRF1 were provided by MRC-PPU and HIS-Titin was provided by Prof. Olga Mayans (University of Konstanz). Titin was cloned into a pMEX3Cb vector to express MBP-Titin by using RF Cloning protocol [53]. Anti-MBP (E8038S, 1:20,000) from New England Biolabs. Anti-6xHis (631212, 1:10,000) from Clontech. Anti-Ubiquitin (646302, 1:1000) from Biolegend. Anti-K48-Specific Ubiquitin (05-1307, 1:1000) and anti-K63-Specific Ubiquitin (05-1313, 1:1000) from Merck-Millipore. Anti-Desmin D93F5 (5332, 1:1000) from Cell Signalling Technology. Anti-MYLPF (MF-5, 1:1000) from Developmental Studies Hybridoma Bank.

### Protein Expression and Purification

Plasmids were transformed into BL21 competent E. coli cells. A colony of each was selected and inoculated in antibiotic-treated LB buffer (For MuRF1 expression LB was treated with 200 mM ZnSO4, prior to IPTG induction) expanding to 2000 mL at 37 °C, grown to OD 600 of 0.6 before adding 250 µM IPTG and left to express overnight at 18 °C. Cultures were pelleted and resuspended in lysis buffer (HIS/HIS-SUMO tag: 50 mM Tris-HCl pH 8.0, 150 mM NaCL, 50 mM Imidazole, 0.5 mM TCEP, 1 mM PMSF. MBP tag: 50 mM Tris-HCl pH 7.5, 150 mM NaCl, 5% Glycerol, 1 mM TCEP, 1 mM PMSF) and then lysed using Emulsiflex C3 Cell Disruptor (Avestin Europe, Mannheim, Germany). Recombinant proteins were purified using a HIS-Trap (GE Healthcare) or Amylose resin (New England Biosciences) as per manufacturer’s instructions. Protein samples were concentrated using 50K centrifugal filters (Amicon, Merck) and stored at -80 °C.

### *In-vitro* Ubiquitylation Assays

The reaction mixture (50 µL) contained 50 mM HEPES pH 7.5, 1 mM DTT, 5 µM ubiquitin, 34 nM HIS-UBE1, 7 µM UBE2, 230 nM MBP-MuRF1 or HIS-SUMO-MuRF1. For experiments involving substrates (i.e., Titin fragment (A160-A170), Desmin or MYLPF) the addition of 1µg of substrate was included in reactions. Reactions were incubated for 1 hr at 37 °C and terminated by the addition of LDS Sample Buffer (Thermo Scientific) to final concentration 1x LDS buffer with 1.25% β-mercaptoethanol. Samples were left overnight to denature before running SDS-page.

### Western Blot

Samples were loaded into 10, 12% or gradient commercial (Invitrogen) Bis-Tris gels for electrophoresis. Gels were run using MOPS buffer (50mM MOPS, 50mM TRIS-Base, 3.465mM SDS, 1.026mM EDTA), 10mins at 100V followed by approx. 60mins at 140V. Proteins were then transferred to PVDF membrane for 1 hr at 100V. Membranes were blocked in 5% skimmed milk powder (Marvel, UK) diluted in Tris-Buffered saline Tween-20 (TBS-T): 137 mM of sodium chloride, 20mM of Tris-Base 7.5 pH, 0.1% of tween-20 for 1 hr and incubated overnight with the requisite primary antibody overnight at 4 °C. Primary antibodies were diluted in 3% BSA in TBS-T. Membranes were washed three times in TBS-T before being incubated in horseradish peroxidase (HRP) conjugated secondary antibodies at room temperature for 1 hr. Membranes were washed three more times in TBS-T before antibody detection using enhanced chemiluminescence horseradish peroxidase substrate detection kit (Millipore, Hertfordshire, UK). Imaging was conducted using a G:Box Chemi-XR5 (Syngene, Cambridgeshire, UK).

### Animal Studies

Male mice from the C57BL/6 strain were raised to 3-4 months of age before denervation as per previous study [38]. Briefly, targeted denervation of the lower limb muscles in the right leg was accomplished through transection of the sciatic nerve. Under isoflurane anaesthesia (2-4% inhalation) and with the use of aseptic surgical techniques, the sciatic nerve was isolated in the mid-thigh region and cut with sharp scissors. Mice were given an analgesic (buprenorphine, 0.1 mg/kg) immediately after the surgery and returned to their cage following recovery. Upon completion of the appropriate time, mice were anaesthetised with isoflurane, and muscles were excised, weighed, frozen in liquid nitrogen, and stored at −80 °C for biochemical analysis. Muscles were also collected from a group of control mice. Once tissue removal was complete, mice were euthanized by exsanguination. All animal procedures were approved by the Institutional Animal Care and Use Committee of the University of Iowa.

### RNA Isolation and qPCR

RNA was isolated from frozen gastrocnemius muscle powder using RNAzol RT reagent (Sigma-Aldrich, St Louis, MO) in accordance with the manufacturer’s instructions. cDNA was synthesised using the iScript Reverse Transcription Supermix kit (BioRad, Hercules, CA) from 1 µg of total RNA. PCR reactions (10 µL) were set up as: 2 µL of cDNA, 0.5 µL (10 µM stock) forward and reverse primers, 5 µL of Power SYBR Green master mix (Thermo Fisher Scientific) and 2 µL of RNA/DNA free water. Gene expression analysis was then performed by quantitative PCR on a Quantstudio 6 Flex Real-time PCR System (Applied Biosystems, Foster City, CA). PCR cycling: hold at 50 °C for 5 min, 10 min hold at 95 °C, before 40 PCR cycles of 95 °C for 15 s followed by 59 °C for 30 s and 72 °C for 30 s. Melt curve analysis at the end of the PCR cycling protocol yielded a single peak. As a result of reference gene instability, gene expression was normalised to tissue weight and subsequently reported as the fold change relative to control muscles, as described previously [54,55]. The mouse primers used in this study are shown in Table S2.

### Statistical Analysis

Data presented as ± SEM. The statistical analyses were performed using Prism (GraphPad Software). one-way ANOVA was performed with Tukey’s post hoc test Values of *P* < 0.05 were considered statistically significant.

## Abbreviations

MuRF1: Muscle-specific RING finger protein 1
RING: Really Interesting New Gene
MYLPF: Myosin Light Chain, Phosphorylatable, Fast Skeletal Muscle Protein
TRIM: Tripartite Motif-Containing
SUMO: Small Ubiquitin-like Modifier
SPR: Surface Plasmon Resonance
MBP: Maltose Binding Protein
ELISA: enzyme-linked immunosorbent assay
CHIP: C-terminus of Hsc70 Interacting Protein
MAFbx: muscle atrophy F-box
MUSA1: muscle ubiquitin ligase of SCF complex in atrophy-1
UBE1: E1 ubiquitin-activating enzyme
UBE2: E2 ubiquitin-conjugating enzyme
TBS-T: Tris-buffered saline Tween-20.

## AUTHOR CONTRIBUTIONS

P.W.J.D. and Y.L. conceived and designed research;

P.W.J.D., L.M.B., D.C.H., and Y.L. performed experiments;

P.W.J.D., L.M.B., and Y.L. prepared figures;

P.W.J.D., and L.M.B. analysed Data;

P.W.J.D, and Y.L. interpreted results of experiments;

P.W.J.D. drafted manuscript;

P.W.J.D. and Y.L. edited and revised manuscript;

P.W.J.D, L.M.B., D.C.H., S.C.B., T.J.K, P.S, and Y.L. approved final version of the manuscript.

## ACKNOWLEDGEMENT

Y.L. was supported by MRC Versus Arthritis Centre for Musculoskeletal Ageing Research (MR/P021220/1). P.D. was supported by MRC Versus Arthritis Centre for Musculoskeletal Ageing Research.

## CONFLICT OF INTEREST

The authors have no conflicts of interest to declare.

## SUPPLEMENTARY TABLES

**Table S1.** UBE2s excluded from *In-vitro* screening.

| Potential UBE2 | Reason for exclusion |
| --- | --- |
| UBE2F | NEDD8-conjugating E2 |
| UBE2I | SUMO-Conjugating E2 |
| UBE2L6 | ISG15-Conjugating E2 |
| UBE2M | NEDD9-Conjugating E2 |
| UBE2QL | Only a probable E2 |
| BIRC6 | Chimeric E2/E3 |
| UBE2U | Only expressed in testis |

**Table S2.**
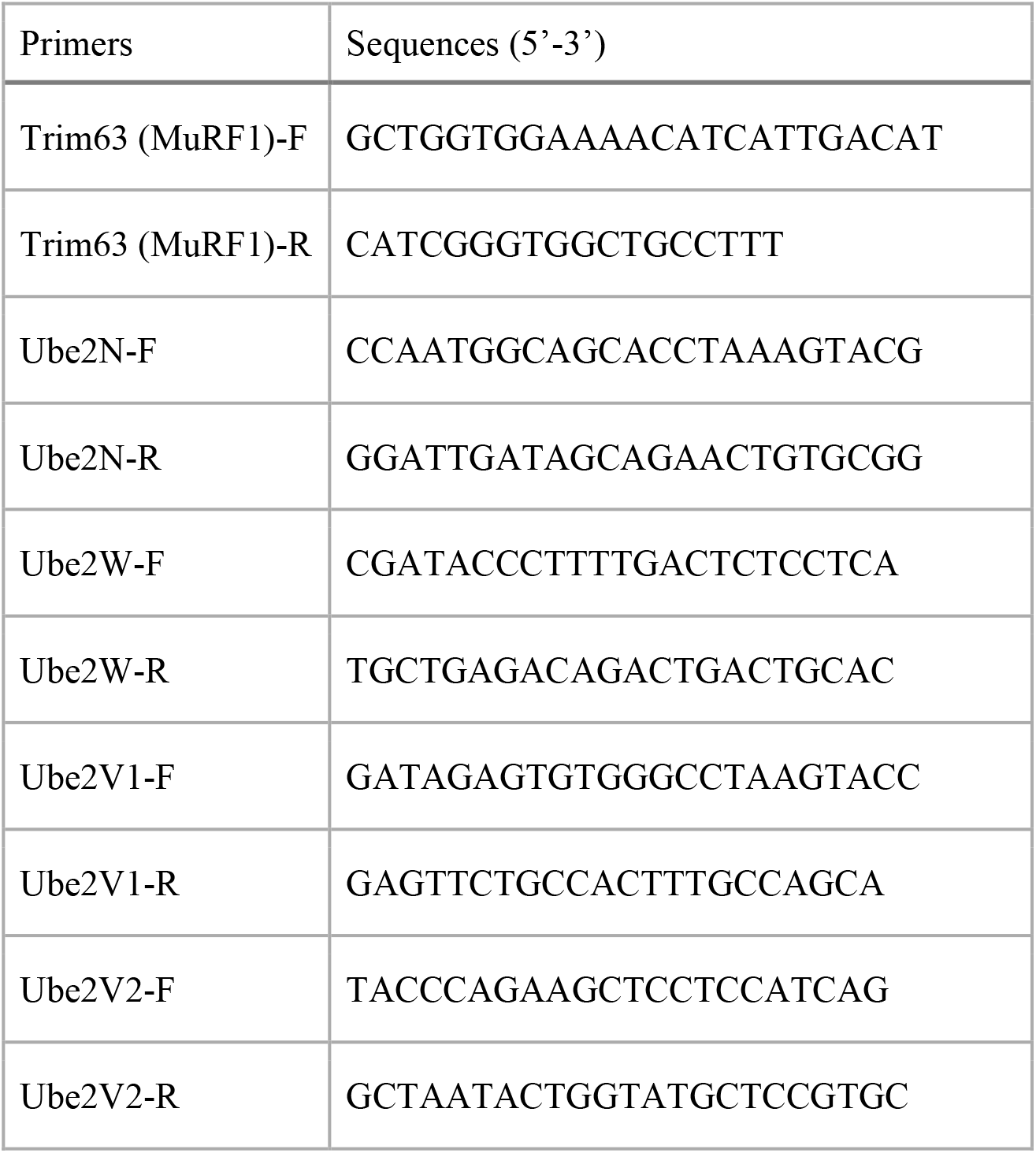
Primers used for qPCR.

## Notes

### Competing Interest Statement

The authors have declared no competing interest.

